# A paleogenetic perspective of the Sabana de Bogotá (Northern South America) population history over the Holocene (9000 – 550 cal BP)

**DOI:** 10.1101/2020.01.24.918425

**Authors:** Miguel Delgado, Freddy Rodríguez, Kalina Kassadjikova, Lars Fehren-Schmitz

**Affiliations:** Consejo Nacional de Investigaciones Científicas y Técnicas (CONICET), Buenos Aires, Argentina; División Antropología, Facultad de Ciencias Naturales y Museo, Universidad Nacional de La Plata, La Plata, Argentina; Ministry of Education Key Laboratory of Contemporary Anthropology Collaborative Innovation Center of Genetics and Development, School of Life Sciences and Human Phenome Institute Fudan University, Shanghai, China; Facultad de Estudios del Patrimonio Cultural, Programa de Arqueología, Universidad Externado de Colombia, Bogotá, Colombia; Anthropology Department & UCSC Human Paleogenomics Lab, University of California Santa Cruz, USA

**Keywords:** Sabana de Bogotá, paleogenetic evidence, population history, Holocene

## Abstract

On the basis of distinct lines of evidence, detailed reconstructions of the Holocene population history of the Sabana de Bogotá (SB) region, Northern South America, have been performed. Currently, there exist two competing models that support temporal continuity or, alternatively, divergence. Despite recent research that lends support to the population discontinuity model, several discrepancies remain, calling for other kinds of evidences to be explored for a more detailed picture of Holocene biocultural evolution. In this study, we analyze the mitochondrial genetic diversity of 30 individuals (including 15 newly reported complete mitochondrial genomes) recovered from several archaeological sites spanning from the late Pleistocene (12,164 cal BP) until the final late Holocene (2,751 cal BP) along with published data from the region dating ∼9,000-550 cal BP in order to investigate diachronic genetic change. Genetic diversity and distance indices were calculated, and demographic models tested in an approximate Bayesian computation (ABC) framework to evaluate whether patterns of genetic affinities of the SB prehispanic populations support genetic continuity or discontinuity. The results show that mitochondrial genomes of the complete dataset fall within the Native American haplogroups A2, B2, C1b, D1 and D4h3a. Haplotype and nucleotide diversity declined over time with further evidence of genetic drift and remarkable reduction of genetic diversity during the final late Holocene. Inter-population distances and the exact test of population differentiation, as well as demographic simulations show no population differentiation and population continuity over time. Consequently, based on the analyzed data, we cannot reject the genetic continuity in the SB region as a plausible population history scenario. However, the restriction of the analyses to the Hyper Variable Region 1 of the mitochondrial genome, and the very low sample size both constitute significant limitations to infer evolutionary history.

## 1. Introduction

The Sabana de Bogotá (SB) in the eastern highlands of Colombia, Northern South America, is a well-known archaeological region that played an important role in the initial human expansion into South America (Correal, 1986; Ardila, 1991; Dillehay, 2000; Aceituno et al., 2013; Delgado et al., 2015ab; Delgado, 2017). In this region, archaeological investigations have recovered abundant lithic, paleobotanical and zooarchaeological evidences in addition to hundreds of burials with well-preserved human skeletal remains dating from 12,164 to 300 cal BP (Correal and van der Hammen, 1977; Correal, 1979, 1990; Ardila, 1984; Botiva, 1989; Correal., 1987; Boada, 1987; Enciso, 1990-1991; Groot, 1992; Orrantia, 1997; Archila and Langebaek, 2015; Triana, 2019 Archila et al., 2020; Ospina and Archila, 2020). In addition, detailed reconstructions of the predominant environmental conditions during the last 18,000 years have been performed on the basis of palynological, glaciomorphological, isotopic and diatom evidences (van der Hammen, 1974; van der Hammen and Hooghiemstra, 1995; Marchant et al., 2002; Vélez et al., 2006; Gómez et al., 2007). Accordingly, this region provides profuse archaeological and paleoecological evidences to perform comprehensive reconstructions of the biological and cultural history of the humans who arrived and settled the region in the late Pleistocene and who evolved throughout the Holocene.

The population history of the SB region has been interpreted both as a process of *in situ* evolution (i.e., the continuous transmission of biological and cultural traits over a defined unit of time) and as a process of biocultural discontinuities (i.e., the cessation of transmission of those traits and eventually their replacement by others). Currently, there exist two competing models supporting population continuity and divergence, respectively termed *Local Evolution Model* (LEM) and *Population Discontinuity Model* (PDM) (*sensu* Delgado, 2012)

### 1.1 Population continuity and discontinuity models

Since the late 1970s, some authors have suggested that specialized hunter-gatherers using the Abriense lithic industry or “edge-trimmed tool tradition” entered the region during the late Pleistocene (*∼*15,000 −16,000 cal BP) and remained there without major population, technological, or economical shifts (excepting the entry *c*. 13,000 cal BP and early disappearance *c*. 11,800 cal BP of a distinct lithic technology named Tequendamiense), until the arrival of agriculturalists during the late Holocene (*c*. 3,400 cal BP) (Correal and van der Hammen, 1977; Hurt et al., 1977; van der Hammen et al., 1990; Correal, 1990) or even until the arrival of European invaders during the 16th century (Rodríguez, 2001, 2007; Rodríguez and and Vargas, 2010). Based on the analysis of traditional craniometric and odontometric traits, some investigators have supported different variants of the continuity scenario. For instance, Rodríguez J.V. and colleagues (Rodríguez, 2001, 2007; Rodríguez and Vargas, 2010) indicated that the morphological patterns representing the same lineage undergo a progressive and subtle transformation throughout the Holocene. In their view, neither the arrival of foreign populations(s) nor changes in the subsistence system or environmental conditions prompted major craniofacial transformations/adaptations. From a slightly different perspective, Neves et al. (2007) suggested that an early population with Paleoamerican cranial morphology entered and evolved locally without important phenotypic transformations and by the initial late Holocene (*c*. 3,400 cal BP) were completely replaced by foreign agricultural populations with an Amerindian cranial pattern. This hypothesis was named the *Local Evolution Model* (LEM) (Delgado, 2012).

On the basis of distinct and independent lines of evidence, some authors have given support to the biocultural discontinuity scenario. Dillehay (2000) detected important cultural changes during the late Pleistocene and early Holocene (*∼*16,000–9,000 cal BP), including technological simplification and economic diversification apparently attributable to the entry of distinct groups. Nieuwenhuis (2002) identified a series of significant shifts in the lithic technology incompatible with the alleged existence of just one lithic techno-complex: i) appearance of more complex artifacts; ii) employment of foreign raw materials; iii) use of tools in non-specialized contexts and activities (i.e. broad spectrum economies); and iv) increasing importance of wood-working and vegetable resources. More recently, Mutillo and colleagues (2017) criticized the chronological and technological basis of the local late Pleistocene/early Holocene lithic techno-complexes (i.e. Abriense and Tequendamiense) that disputes the alleged homogeneity of the main indicator of population continuity, such as the persistence of the Abriense kind over the last 12,000 years. Additionally, some authors have suggested that the existence of intersocietal interactions and widespread exchange networks between the SB and other regions may have promoted cultural change, that is, induced cultural innovations and/or modifications (Correal and van der Hammen, 1977; Nieuwenhuis, 2002; Aceituno and Loaiza, 2007; López, 2008). In fact, Gnecco (1999, 2003) and Nieuwenhuis (2002) suggest that the foreign lithic raw materials (chert from the Middle Magdalena Valley) found in El Abra, Tequendama and Tibitó can indicate both territorial mobility and intergroup contacts during the late Pleistocene/early Holocene, which could have promoted gene flow. Cárdenas (2002) and Delgado (2018), using biochemical evidences (stable isotopes), criticized the alleged specialized economies, stressed the presence of early and mid-Holocene hunter-gatherers with mixed C_3_/C_4_ diets that may indicate the entry of foreign lowland groups, and suggested that the emergence of tropical crop agriculture during the initial late Holocene (*c*. 3,700 cal BP) is related to population dispersals rather than an *in situ* development. Finally, some authors addressing the skeletal diversity at the regional level have detected significant changes in the craniofacial and dental morphology probably associated with adaptations to environmental and subsistence shifts as well as with population contacts, retractions/extinctions, and replacements during the middle (*c*. 5,900-4,800 cal BP) and final late Holocene (*c*. 2,700 cal BP) (Ardila, 1984; Correal, 1990; Cárdenas, 2002; Delgado, 2012, 2015, 2017, 2018; Rodríguez Flórez and Colantonio, 2015). This scenario was named *Population Discontinuity Model* (PDM) (Delgado, 2012).

Notwithstanding the support for one of the existing population history models, several gaps remain in Holocene human history that are very difficult to address using the available archaeological, bioarchaeological and paleoecological data. This suggests that other lines of evidence must be explored. Accordingly, the aim of the present study is to use ancient DNA to further evaluate if the genetic affinities patterns of the pre-Hispanic populations inhabiting the SB region during the last ∼12,000 years support one of the two regional population history models (LEM or PDM).

### 1.2 The area of study

The SB is a high plain located 2,600 m above sea level (asl) on the eastern side of the Andean cordillera in Northwestern South America (Figure 1). This high plain has an approximate area of 1,400 km^2^, with slopes ranging between 3,000 and 4,000 m asl in the north and south side respectively (van der Hammen and González, 1960; Dueñas, 1980). In the region, the most important river is the Bogotá, although some lagoons also exist. On the western side of the SB lies the Magdalena River Valley and on the eastern side, the mountains slope down to the eastern plains. Geologically, the SB is a Tertiary basin product of a geological-evolutionary process whose last event was the raising of the basal sediments of the Tilatá formation at the end of the Pliocene (van der Hammen and González, 1960; Dueñas, 1980). Throughout the Pleistocene and the Holocene, lake clays were deposited from the Sabana formation, which filled the basin and contemporary landscape of the SB (Dueñas, 1980; van der Hammen and Correal, 1992). At the present time this region supports an Andean forest characterized by the presence of *Weinmannia* sp. div., *Quercus*, and other species (van der Hammen and González, 1960). There are no seasonal temperature fluctuations, only differences in rainfall, with a wet season twice a year. The annual temperature varies between 12 and 18°C and the annual rainfall varies between 500 and 1,500 mm (van der Hammen and Correal, 1992). Regarding the faunal composition of the SB, the archaeological evidence suggests a high density of species including mammals (v.g. *Odocoileus virginianus*, *Mazama Sp*, *Cavia porcellus*); reptiles (*Kinosternon postinginale*, *Crocodlylia sp*); fishes (v.g. *Eremophilus mutisii*, *Pygidium bogotense*); birds (v.g. *Familia anatidae*, *Familia ralidae*) and crustaceous-gastropods (v.g. *Neostrengeria macropa*; *Drymaeus gratus*).

**Figure 1.**
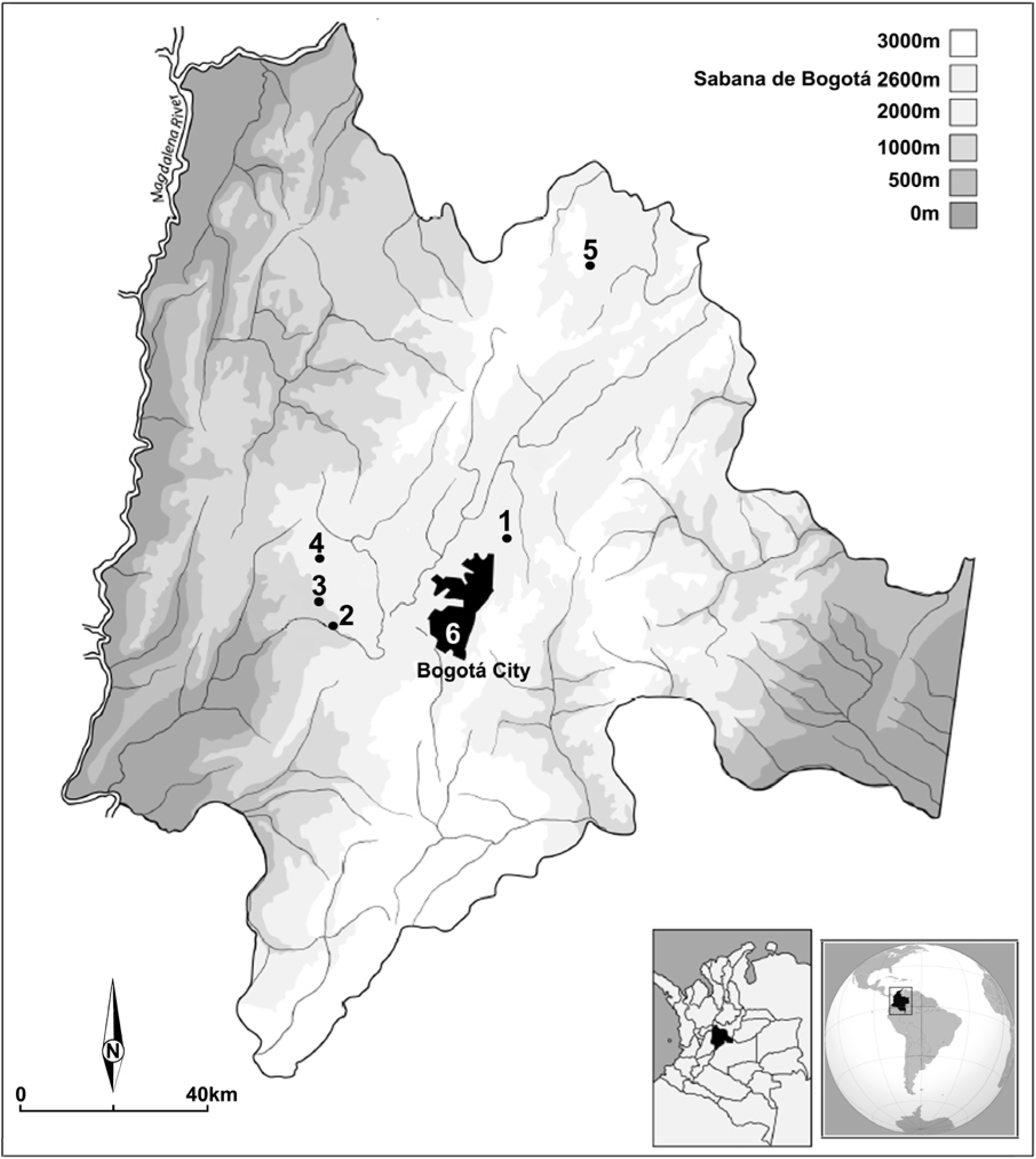
The Sabana de Bogotá region showing the location of archaeological sites with human skeletal remains investigated. 1= Checua; 2= Tequendama; 3= Aguazuque; 4= Vistahermosa; 5= Ubaté; 6 = Tibanica.

## 2. Materials

### 2.1. The human skeletal remains investigated

The SB is one of the few archaeological regions in Northern South America where hundreds of human skeletal remains from the last ∼12,000 years have been recovered. For the present study, 30 individuals (Figure 1; Table 1) representing distinct hunter-gatherer, horticultural and agricultural societies from several open-air and rock shelter multicomponent sites in the SB region were recovered and analyzed. All individuals were in relatively good macroscopic preservation conditions typical of the study area and most of them have reliable chronological contexts. The radiocarbon dates were calibrated using OxCal 4.3 (Bronk Ramsey, 2009) using the SHCal 13 curve (Hogg et al., 2013) at 2-σ (95.4% probability).

**Table 1.**
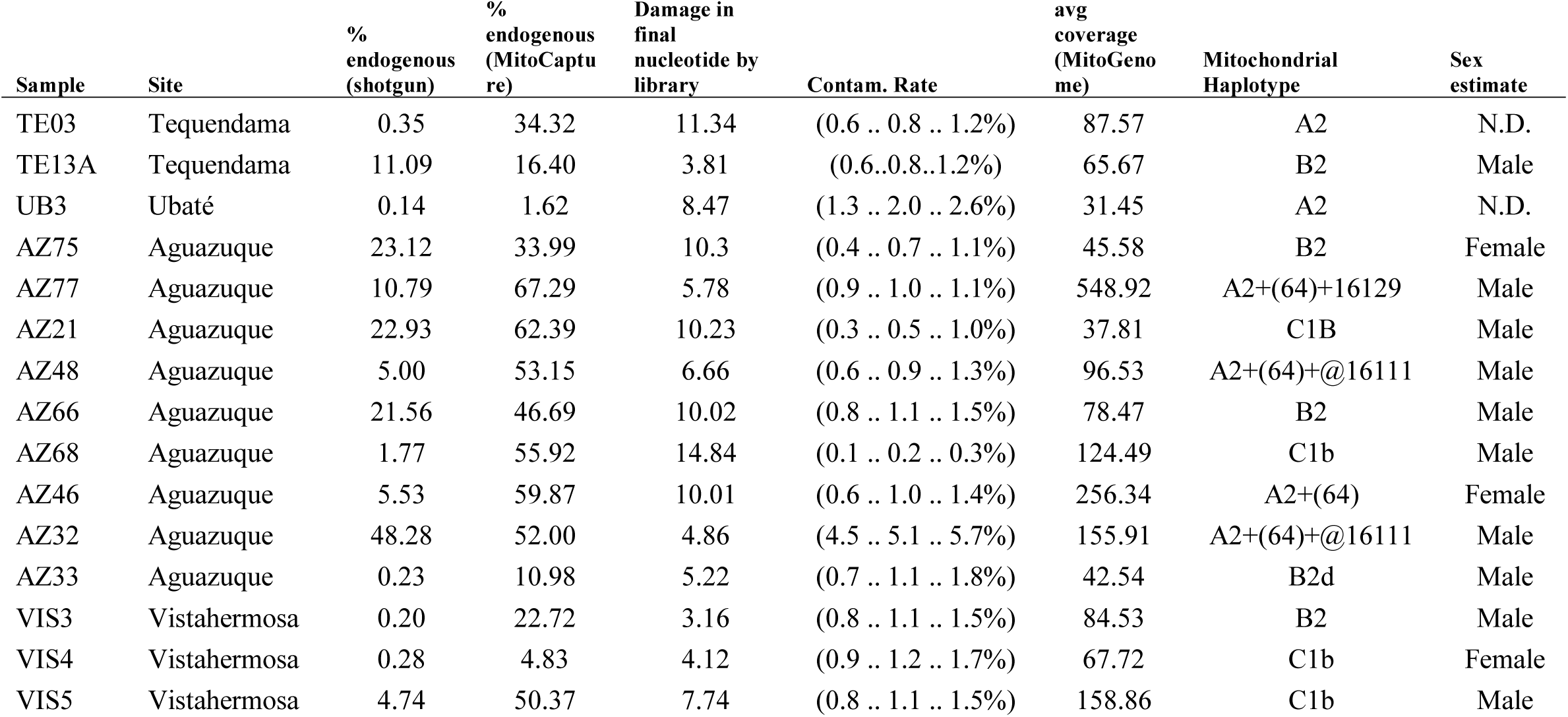
Sequencing statistics and mitochondrial haplotypes for the 15 mitochondrial genomes

| Sample | Site | % endogenous (shotgun) | % endogenous (MitoCapture) | Damage in final nucleotide by library | Contam. Rate | avg coverage (MitoGenome) | Mitochondrial Haplotype | Sex estimate |
| --- | --- | --- | --- | --- | --- | --- | --- | --- |
| TE03 | Tequendama | 0.35 | 34.32 | 11.34 | (0.6 .. 0.8 .. 1.2%) | 87.57 | A2 | N.D. |
| TE13A | Tequendama | 11.09 | 16.40 | 3.81 | (0.6..0.8..1.2%) | 65.67 | B2 | Male |
| UB3 | Ubaté | 0.14 | 1.62 | 8.47 | (1.3 .. 2.0 .. 2.6%) | 31.45 | A2 | N.D. |
| AZ75 | Aguazuque | 23.12 | 33.99 | 10.3 | (0.4 .. 0.7 .. 1.1%) | 45.58 | B2 | Female |
| AZ77 | Aguazuque | 10.79 | 67.29 | 5.78 | (0.9 .. 1.0 .. 1.1%) | 548.92 | A2+(64)+16129 | Male |
| AZ21 | Aguazuque | 22.93 | 62.39 | 10.23 | (0.3 .. 0.5 .. 1.0%) | 37.81 | C1B | Male |
| AZ48 | Aguazuque | 5.00 | 53.15 | 6.66 | (0.6 .. 0.9 .. 1.3%) | 96.53 | A2+(64)+@16111 | Male |
| AZ66 | Aguazuque | 21.56 | 46.69 | 10.02 | (0.8 .. 1.1 .. 1.5%) | 78.47 | B2 | Male |
| AZ68 | Aguazuque | 1.77 | 55.92 | 14.84 | (0.1 .. 0.2 .. 0.3%) | 124.49 | C1b | Male |
| AZ46 | Aguazuque | 5.53 | 59.87 | 10.01 | (0.6 .. 1.0 .. 1.4%) | 256.34 | A2+(64) | Female |
| AZ32 | Aguazuque | 48.28 | 52.00 | 4.86 | (4.5 .. 5.1 .. 5.7%) | 155.91 | A2+(64)+@16111 | Male |
| AZ33 | Aguazuque | 0.23 | 10.98 | 5.22 | (0.7 .. 1.1 .. 1.8%) | 42.54 | B2d | Male |
| VIS3 | Vistahermosa | 0.20 | 22.72 | 3.16 | (0.8 .. 1.1 .. 1.5%) | 84.53 | B2 | Male |
| VIS4 | Vistahermosa | 0.28 | 4.83 | 4.12 | (0.9 .. 1.2 .. 1.7%) | 67.72 | C1b | Female |
| VIS5 | Vistahermosa | 4.74 | 50.37 | 7.74 | (0.8 .. 1.1 .. 1.5%) | 158.86 | C1b | Male |

Seven early hunter-gatherers from Sueva, Galindo, Tequendama and Potreroalto whose chronological range corresponds to the final Pleistocene and early Holocene (∼7,000 and 12,000 cal BP) were investigated. The skeleton from Sueva I rock shelter (2,690 m asl) recovered from the stratum 3 dated at 12,164–11,231 cal BP (10,090±90 ^14^C BP; GrN-8111) corresponds to a female (Correal, 1979). Another male individual was included from the open-air site Galindo I (2,598 m asl), found in level 2 and dated by association to 8,722–8,375 cal BP (7,730±60 ^14^C BP GrN-16345) using charcoal from the burial (Pinto, 2003). Three skeletons (2 males and 1 undetermined) from the multicomponent site Tequendama I and II were also investigated. This rock shelter (2,570 m asl) was occupied from the final Pleistocene 13,581 –11,766 (10,920±260 GrN-6539) until the late Holocene 2,347–2,121 cal BP (2,225±35 ^14^C BP GrN-6536) (Correal and van der Hammen, 1977). The early/middle Holocene burials with human skeletal remains are restricted to strata 7a and 8ab whose chronological range spans from ∼9,000 to 6,500 cal BP (Delgado, 2015). One of the skeletons investigated was dated by conventional methods (i.e., beta counting, Correal and van der Hammen, 1977) and was corrected by isotopic fractionation to 7,162– 6,633 cal BP (burial 13 6,056±51 ^14^C BP GrN-7478) (Delgado, 2015). The other two skeletons from stratum 8 are supposed to be associated with an additional two dates corrected by isotopic fractionation to 8,224–7,935 cal BP (7,269±65 ^14^C BP GrN-7477) and 6,860–6,451 cal BP (5,848±56 ^14^C BP GrN-7476) (Delgado, 2015). Finally, two individuals (a male and a female) from the open-air site Potreroalto (2,610 m asl) were included. Both individuals were dated by AMS to 7,966–7,430 cal BP (5,910±70 ^14^C BP Beta-104490) and 7,000–6,479 (6,830±110 ^14^C BP Beta-104491) respectively (Orrantia, 1997).

The middle Holocene period is represented by fourteen individuals comprising generalized hunter-gatherers, fishermen and early horticulturalists from four open-air and rock shelter sites: Ubaté, Bonacá, Chia III and Aguazuque. Four individuals (1 male, 1 female and 2 undetermined) are from the recently excavated open-air sites Ubaté (2,550 m asl) and Bonaca (2,600 m asl), which provided additional evidences on the mid-Holocene hunter-gatherer way of life (Archila and Langebaek, 2015; Pérez et al., 2016). Such skeletons were dated by AMS to 6,491–6,305 cal BP (5,620±30 ^14^C BP Beta-398660) and 6,295–6,170 (5,400±30 ^14^C BP Beta-398659) for Ubaté (Archila and Langebaek, 2015) and at 6,474–6,276 (5,560±40 ^14^C BP Beta-347992) and (5,540±30 ^14^C BP Beta-347993) for Bonaca (Pérez et al., 2016). One additional female individual was included from the Chia III rock shelter (2,610 m asl) (Ardila, 1984). Despite several burials were uncovered in this site, only two individuals from burial five presented good preservation. One of these individuals was dated by beta counting techniques and corrected by isotopic fractionation to 6,213–5,581 cal BP (5,084±103 ^14^C BP GrN-12122) (Delgado, 2015). Finally, nine individuals came from the multicomponent open-air site Aguazuque (2613 m asl) whose chronological range spans from 5,919–5,650 (5,069±47 ^14^C BP GrN-14477) to 2,947–2,751 cal BP (2,769±45 ^14^C BP GrN-14479) (Correal, 1990). This site is key for understanding SB human-history because it is one of the few existing sites that present evidence on the transition between hunter-gatherers, horticulturalists and agriculturalists. All radiocarbon dates derived from human bone and were therefore corrected by isotopic fractionation (Delgado, 2015). The mid-Holocene individuals investigated were from units 3 and 4, corresponding to the first occupation whose chronological range is from 5,941–5,650 (5,069±47 ^14^C BP GrN-14477) to 4,821–4,421 cal BP (4,074±41 ^14^C BP GrN-12930) (Correal, 1990; Delgado, 2015).

Lastly, ten individuals (2 females, 6 males and 2 undetermined) from two open-air sites Aguazuque and Vistahermosa representing the first horticultural and agricultural populations from the initial late Holocene period were investigated. Four individuals are from the third occupation of Aguazuque corresponding to the unit 4.2 dated to 4,516–4,470 (corrected by isotopic fractionation 3,894±42 ^14^C BP GrN-14478) and two are from the fifth occupation corresponding to the unit 5.2 dated to 3,057–3,049 (2,769±45 ^14^C BP GrN-14479 corrected by isotopic fractionation) (Correal, 1990; Delgado, 2015). Four individuals from the open-air site Vistahermosa (2,590 m asl) were included, who were recovered from strata 1 and 2 (Correal, 1987). The stratum 1 was dated using charcoal associate with the burials to 3,833–3,558 cal BP (3,135±35 ^14^C BP GrN-12928), while the stratum 2 was dated to 3,453–3,210 cal BP (3,410±35 ^14^C BP GrN-12929).

### 2.2 Comparative data

In order to obtain a more detailed picture on the diachronic genetic differentiation at the SB region, some published sequences corresponding to early/middle and final late Holocene individuals were collated from the literature. Unfortunately, few reliable comparative studies exist, which mostly investigate only the HVI and HVII regions of the mtDNA genome. For the early to initial late Holocene periods, 12 individuals from the Checua site were included (Díaz-Matallana, 2015). Distinct to such study, the individuals included were not considered belonging to the early Holocene but were distributed into three distinct chronological groups, taking into account clear stratigraphic and chronological differences reported in the original study (Groot, 1992): 1) n=4 (CHI3, CHI1, CH08, CH07) early Holocene (8,868–8,708 cal BP; 7,154–7,117 cal BP; 2) n=6 (CH01, CH02, CHII03, CHII04, CHII05, CHII06) middle Holocene (5,914–5,579 cal BP) and 3) n=2 (CHI03a, CHI03b) initial late Holocene (3,594–3,141 cal BP; 2,997–2,486 cal BP). Finally, two samples from the final late Holocene period were included, one corresponding to the late Muisca period (Pérez L, 2015) and the other attributed to the Guane archaeological entity from the neighbor Santander province (Casas-Vargas et al., 2011). The Muisca sample was integrated, after considering several quality controls, by 8 individuals out of 74 from the Tibanica site investigated by Pérez L (2015) whose chronological range is between 935–744 cal BP and 651–540 cal BP (maximum and minimum range of 16 AMS dates) (Miller, 2016). The Guane sample is integrated by 14 individuals from the La Purnia site (Santander Province) dated to 1,185–732 cal BP (Casas-Vargas et al., 2011). The inclusion of the Guane sample from a neighbor region, despite the small portion of the HVR1 sequenced, is justified given the underrepresentation of samples from the final late Holocene period and because the Guane populations closely interacted with Muisca groups in terms of economy, exchange, and social relationships according to ethnohistoric accounts (Pérez P, 2001).

## 3. Methods

All samples in this study were processed in the dedicated clean rooms at UCSC Paleogenomics in Santa Cruz (USA), following strict procedures to minimize contamination (Llamas et al., 2017). DNA was extracted from bone or tooth samples using a method that is optimized to retain small DNA fragments (Dabney et al., 2013; Boessenkool et al., 2016). Single-stranded sequencing libraries were prepared from all DNA extracts using the Santa Cruz methods described by Troll et al. (Troll et al., 2019). DNA extracts were treated with uracil-DNA glycosylase (UDG) to greatly reduce the presence of errors characteristic of ancient DNA at all sites except for the terminal nucleotides (Rohland et al., 2014). The preservation and endogenous DNA content of the libraries was then tested by sequencing on an Illumina MiSeq platform at UCPL (UCSC Paleogenomics Labs), using 2×75 bp paired end sequencing (∼300k reads per library).

To screen sequence data computationally, we merged paired reads that overlapped by at least 15 nucleotides using SeqPrep (https://github.com/jstjohn/SeqPrep), taking the highest quality base to represent each nucleotide, and then mapped the sequences to the human genome reference sequence (GRCh37 from the 1000 Genomes project) using the *samse* command of the Burrows-Wheeler Aligner (BWA) (version 0.6.1) (Li and Durbin, 2009). Libraries showing suitable content of endogenous DNA fragments (%endogenous > 1%) and high complexity (less than 0.5% duplication after 300k reads) were then enriched for mitochondrial DNA using the myBaits Mito Human Global Panel in-solution hybridization Kit from Arbor Biosciences (Ann Arbor, MI). The libraries were captured following the manufacturer’s instructions (http://www.mycroarray.com/pdf/MYbaits-manual-v3.pdf). The captured libraries were amplified for 20 cycles with IS4 and indexed P7 primers and subsequently sequenced on an Illumina NextSeq 500 with ∼1000k reads for each library.

The raw data sequenced from the enriched libraries was analyzed as described above. To test the authenticity of the sequencing data, we measured the rate of damage in the first nucleotide using MapDamage 2.0 (Jonsson et al., 2013), flagging individuals as potentially contaminated if they had a less than 3% cytosine-to-thymine substitution rate in the first nucleotide. We further estimated mitochondrial contamination rates employing the modules contDeam and mtCont implemented in the software tool SCHMUTZI using the recommended parameters (Renaud et al., 2015). Chromosomal sex was determined as described in Skoglund et al. (2013). To identify the mitochondrial haplotypes of the individuals, we manually analyzed each variant, as described in Llamas et al. (2016). The assembly and the resulting list of SNPs were verified manually and compared to SNPs reported at phylotree.org (mtDNA tree Build 17 [18 Feb 2016]) (Van Oven, 2015). Following recommendations in van Oven and Kayser 2009 (Van Oven and Kayser, 2009), we excluded common indels and mutation hotspots at nucleotide positions 309.1C(C), 315.1C, AC indels at 515–522, 16182C, 16183C, 16193.1C(C), and C16519T. We embedded the consensus mitochondrial genomes in the existing mitochondrial tree (mtDNA tree Build 17 [18 Feb 2016]) using the online tool HaploGrep2 (Weissensteiner et al., 2016) to determine the haplotypes.

Due to the lack of ancient whole mitochondrial genomes from the study region, the obtained data was then merged with a dataset of 20 Early to Final Late Holocene HVR1 sequences from the Sabana de Bogotá region (Díaz-Matallana, 2015; Pérez L, 2015; Casas-Vargas et al., 2011) (Supplementary Table 1 and 2) and ∼1,750 modern mitochondrial HVR1 sequences grouped into 42 populations from Central and South America (Supplementary Table 3). For all subsequent population genetic analysis, the data set was restricted to 345 nucleotide positions of the mitochondrial HVR1 region (nucleotide position 16,056-16,400, relative to the revised Cambridge reference sequence (rCRS) Andrews et al. (1999). The ancient mitochondrial data obtained in this study was grouped with the other ancient mitochondrial HVR1 sequences into four broader chronological periods to increase sample size in each group: Early Holocene (∼9,000–7,000 cal BP,n=6), Middle Holocene (∼6,500–4,500 cal BP, n=14), Initial Late Holocene (∼3,600–2,700 cal BP; N=7), Final Late Holocene (∼950–550 cal BP; n=8) (see Table 2; Supplementary Table 1 and 2).

**Table 2.** Early- to Late Holocene HVR1 sequences integrated in this study and chronological grouping

| Sample Id | Archaeological Site | Chronology (14C BP) | Period / Chronological Group | Economy | Haplogroup | Reference |
| --- | --- | --- | --- | --- | --- | --- |
| CHI-11 | Checua | 7800-6000 | Early Holocene | Hunter-Gatherers | A2 | Diaz-Matallana 2015 |
| CHI-13 | Checua | 7800-6000 | Early Holocene | Hunter-Gatherers | A2 | Diaz-Matallana 2015 |
| CHI-07 | Checua | 7800-6000 | Early Holocene | Hunter-Gatherers | B2 | Diaz-Matallana 2015 |
| CHI-08 | Checua | 7800-6000 | Early Holocene | Hunter-Gatherers | C1 | Diaz-Matallana 2015 |
| TE03 | Tequendama | 7000-6000 | Early Holocene | Hunter-Gatherers | A2+(64) | this study |
| TE13 | Tequendama | 7000-6000 | Early Holocene | Hunter-Gatherers | B2 | this study |
| CHII-03 | Checua | 5021 ± 202 * | Middle Holocene | Hunter-Gatherers | A2 | Diaz-Matallana 2015 |
| CHII-04 | Checua | 5021 ± 202 * | Middle Holocene | Hunter-Gatherers | A2 | Diaz-Matallana 2015 |
| CHII-05 | Checua | 5021 ± 202 * | Middle Holocene | Hunter-Gatherers | A2 | Diaz-Matallana 2015 |
| CHII-06 | Checua | 5021 ± 202 * | Middle Holocene | Hunter-Gatherers | B2 | Diaz-Matallana 2015 |
| CHI-1 | Checua | 6000-5000 | Middle Holocene | Hunter-Gatherers | B2 | Diaz-Matallana 2015 |
| CHI-02 | Checua | 6000-5000 | Middle Holocene | Hunter-Gatherers | D4h3a | Diaz-Matallana 2015 |
| AZ77 | Aguazuque | 5025 ± 40 | Middle Holocene | Hunter-Gatherers | A2+(64)+16129 | this study |
| AZ21 | Aguazuque | 4030 ± 35 | Middle Holocene | Hunter-Gatherers | C1b | this study |
| AZ46 | Aguazuque | 3850 ± 35 | Middle Holocene | Hunter-Gatherers | A2+(64) | this study |
| AZ48 | Aguazuque | 3850 ± 35 | Middle Holocene | Hunter-Gatherers | A2+(64)+@16111 | this study |
| AZ66 | Aguazuque | 4030 ± 35 | Middle Holocene | Hunter-Gatherers | B2 | this study |
| AZ75 | Aguazuque | 5025 ± 40 | Middle Holocene | Hunter-Gatherers | B2 | this study |
| UB3 | Ubaté | 5620 ± 30 | Middle Holocene | Hunter-Gatherers | A2+(64) | this study |
| AZ68 | Aguazuque | 4030 ± 35 | Middle Holocene | Hunter-Gatherers | C1b | this study |
| CHI-03a | Checua | 3200-2700 | Initial late Holocene | Hunter-Gatherers | A2 | Diaz-Matallana 2015 |
| CHI-03b | Checua | 3200-2700 | Initial late Holocene | Hunter-Gatherers | A2 | Diaz-Matallana 2015 |
| AZ32 | Aguazuque | 2725 ± 35 | Initial late Holocene | Horticulturalists | A2+(64)+@16111 | this study |
| AZ33 | Aguazuque | 2725 ± 35 | Initial late Holocene | Horticulturalists | B2d | this study |
| VIS3 | Vista Hermosa | 3410 ± 35 - 3135 ± 35 | Initial late Holocene | Hunter-Gatherers | B2 | this study |
| VIS4 | Vista Hermosa | 3410 ± 35 - 3135 ± 35 | Initial late Holocene | Hunter-Gatherers | C1b | this study |
| VIS5 | Vista Hermosa | 3410 ± 35 - 3135 ± 35 | Initial late Holocene | Hunter-Gatherers | C1b | this study |
| M803 | Tibanica | 990 ± 30 - 640 ± 20 ** | Final late Holocene | Agriculturalists | D1 | Perez 2015 |
| M816 | Tibanica | 990 ± 30 - 640 ± 20 ** | Final late Holocene | Agriculturalists | A2+(64)+@16111 | Perez 2015 |
| M841 | Tibanica | 990 ± 30 - 640 ± 20 ** | Final late Holocene | Agriculturalists | A2+(64)+@16111 | Perez 2015 |
| M1166 | Tibanica | 990 ± 30 - 640 ± 20 ** | Final late Holocene | Agriculturalists | B2 | Perez 2015 |
| M2208 | Tibanica | 990 ± 30 - 640 ± 20 ** | Final late Holocene | Agriculturalists | A2+(64)+@16111 | Perez 2015 |
| M3168B | Tibanica | 990 ± 30 - 640 ± 20 ** | Final late Holocene | Agriculturalists | A2 | Perez 2015 |
| M3295 | Tibanica | 990 ± 30 - 640 ± 20 ** | Final late Holocene | Agriculturalists | B2 | Perez 2015 |
| M3305 | Tibanica | 990 ± 30 - 640 ± 20 ** | Final late Holocene | Agriculturalists | B2 | Perez 2015 |

\*\*Maximum and minimum range of 16 AMS dates
\* Date obtained from electron paramagnetic resonance on dental enamel

Haplotype diversity (Hd) and nucleotide diversity (π) indices within populations and geographical groups were calculated and biological distances between the populations and groups were estimated from the HVR1 sequence data. Pairwise F_ST_ values, derived Slatkin distances for populations with short divergence times (Slatkin, 1995), and also the diversity indices were calculated using Arlequin 3.5 (Excoffier and Lischer, 2010). Genetic distances between haplotypes and mean distances between groups were calculated employing the Tamura and Nei distance model with gamma correction (Tamura & Nei, 1993) with the suggested gamma value of 0.26 for mitochondrial HVR1 data (Meyer et al., 1999). To further test for population continuity/discontinuity between the ancient Colombian samples, we performed exact tests of population differentiation (Goudet et al., 1996) with Arlequin. All described tests were performed with at least 10,000 permutations.

To test if observed genetic distances between the Early- and Final Late Holocene populations from the Bogota region could be explained by population divergence or reflect genetic drift we tested different demographic scenarios employing the approximate Bayesian computation (ABC) method, implemented in DIYABC v 2.0.1(Cornuet et al., 2014). We tested four simplified divergence models: 1. Genetic continuity through all four chronological groups (Early Holocene – Final Late Holocene); 2. Discontinuity between the Middle Holocene and Initial Late Holocene (both Early- and Middle Holocene and Initial- and Final Late Holocene forming separate branches that diverged before 10,000 BP); 3. Population discontinuity between Initial Late Holocene and Final Late Holocene, with continuity between Early and Initial Late Holocene; 4. Population discontinuity between Middle Holocene and Initial Late Holocene, and between Initial late Holocene and Final Late Holocene. In all scenarios, we assumed that each population maintained a constant size over time and post divergence from the ancestral population. We used TN93+G as the substitution model and the mtDNA mutation rate determined by Soares et al. (2009). Due to the low number of nucleotide position in the data set, we restricted the compared summary statistics to interpopulation F_ST_ values. Each scenario was simulated one million times. Posterior probabilities of the modeled scenarios were estimated using a logistic regression approach (Cornuet et al., 2014) with the 1% of simulated datasets possessing the smallest Euclidian distances to the observed dataset. Goodness-of-fit between the simulated and real datasets was evaluated using principal component analysis in DIYABC.

## 4. Results

Of the 30 samples tested, only 15 libraries showed sufficient DNA preservation—as measured in %endogenous DNA content, and complexity—following the initial shotgun sequencing screening. The 15 libraries were subsequently enriched for mitochondrial DNA and we obtained whole mitochondrial genomes for all of them, with average coverage ranging from ∼20x to ∼550x. Damage profiles and mitochondrial contamination estimates for all 15 sequenced samples indicate authentic ancient DNA (Table 1). The mitochondrial genomes all fall within the Native American haplogroups A2, B2, and C1b. Detailed information on the haplotypes can be found in Table 1, Supplementary Table 1 and 2, and Figure 2. Two individuals, one from the Early Holocene levels of the site Tequendama (TE03), and one from the Middle Holocene site Ubaté (UB3) share the exact A2+(64)+@16111 haplotype (see Supplementary Table 2; Figure 2). The haplogroup is also found in three Final Late Holocene individuals from the late Muisca site Tibanica, however only HVR1 data is available for comparison from that site (Perez L, 2015). As outlined in the methods, the obtained mitogenomes were then grouped with 20 ancient HVR1 sequences from the SB region into four chronological groups (Table 2; Supplementary Table 1). Native American mitochondrial haplogroup frequencies for each group can be found in Table 3. Throughout all four time periods haplogroup A2 is found to be most frequent (43-50%), followed by haplogroup B2 (28-38%). Only five individuals presented the haplogroup C1, and one individual was found to belong into haplogroup D1 and one into D4h3. However, the low overall sample size (n= 35) has to be considered. The observed haplogroup frequencies for each of the four chronological groups roughly resemble haplogroup frequencies observed for modern populations around the Sabana de Bogotá region, and along Northern Colombia, Venezuela and Southern Central America (cf. Bisso-Machado et al., 2012; Melton et al., 2013; Diaz-Matallana et al., 2016). Nucleotide Diversity (π) estimates for all four chronological groups (restricted to HVR1) range between 0.0155 and 0.0105, and Hd between 0.933 and 0.857 (Table 3). Both π and Hd decline from the Early Holocene to the Final Late Holocene. The diversity indices fall well into the range observed in other modern populations from Central and South America. However, at least the Final Late Holocene samples the Hd falls into the range of highly drifted groups with low diversity like several Amazonian groups (Wang et al., 2007). We have to caution that the obtained diversity values might be biased due to the low sample size.

**Figure 2.**
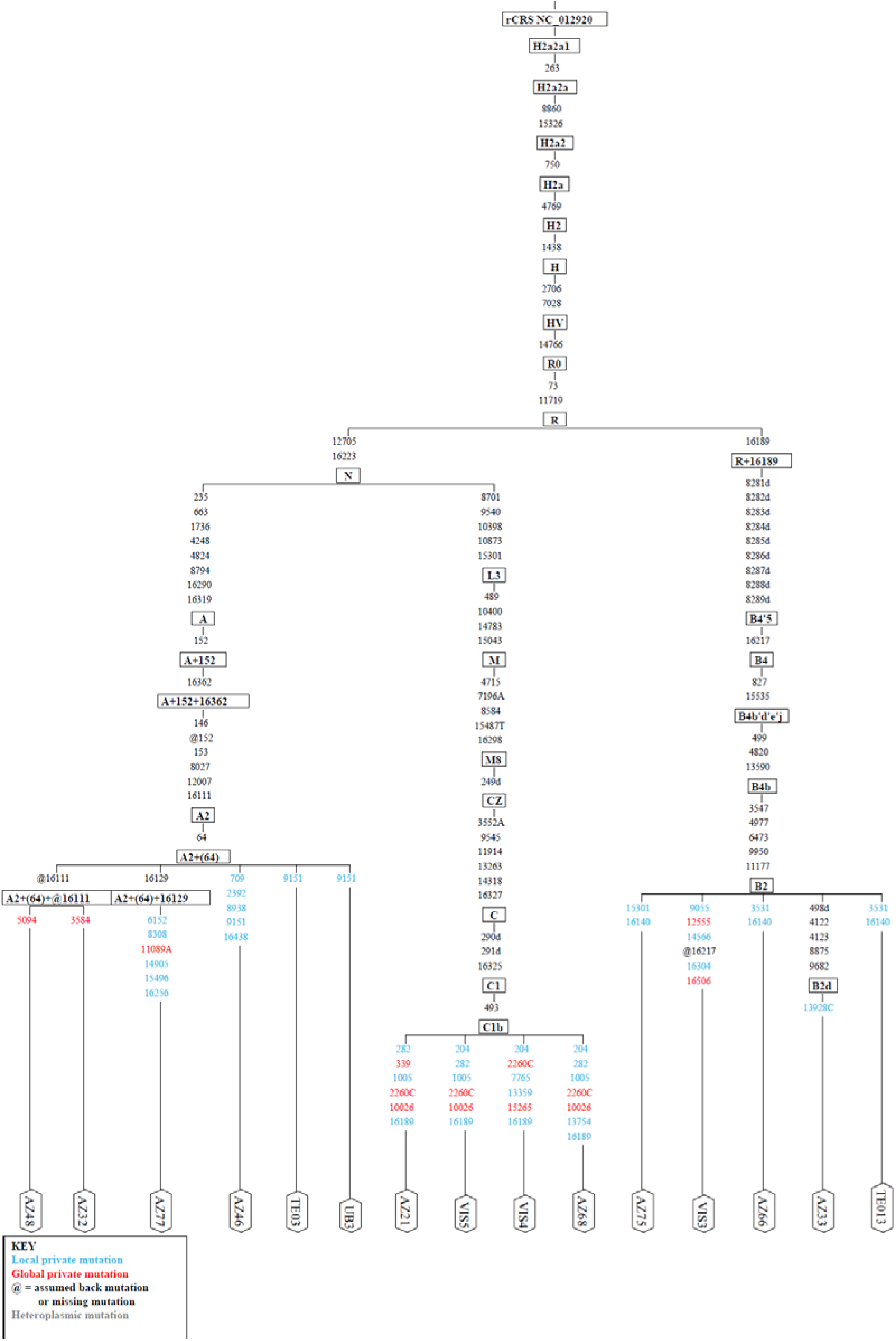
Phylogenetic tree of the 15 new mitochondrial genomes presented in this study, generated by HaploGrep2.

**Table 3.** Diversity statistics and mtDNA haplogroup frequencies for the four chronological groups

|  | **n** | **Haplotypes**<br>**(h)** | **Segregating**<br>**Sites (S)** | **$\pi$** | **Hd** | **mtHaplogroup Frequency** |  |  |  |  |
| --- | --- | --- | --- | --- | --- | --- | --- | --- | --- | --- |
|  |  |  |  |  |  | **A2** | **B2** | **C1** | **D1** | **D4h3** |
| EH | 6 | 6 | 13 | 0.0155 | 0.9333 | 0.50 | 0.33 | 0.17 | 0.00 | 0.00 |
| MH | 13 | 8 | 16 | 0.0134 | 0.8974 | 0.50 | 0.28 | 0.14 | 0.00 | 0.07 |
| ILH | 7 | 5 | 11 | 0.0135 | 0.9048 | 0.43 | 0.29 | 0.29 | 0.00 | 0.00 |
| FLH | 8 | 5 | 9 | 0.0105 | 0.8571 | 0.50 | 0.38 | 0.00 | 0.12 | 0.00 |

To test if the mitochondrial data fits either the postulated LEM or PDM models, we calculated pairwise inter-population distances between the four chronological groups. Each comparison returned negative and non-significant F_ST_-values (F_ST_ = −0.1-0.05, p= 0.88-0.67; Table 4), meaning that we cannot reject the null-hypothesis of no-differentiation (i.e., LEM). The exact test of population differentiation additionally finds no differentiation between the groups (p=1.00). All four chronological groups exhibit zero to low genetic distance to modern Embera and Wounan from Southern Panama (F_ST_= 0-0.016; p=0.42-0.91). The inter-population F_ST_ distances for the whole comparative dataset are visualized with non-metric multidimensional scaling (MDS) in Figure 3. As shown in this graphic (stress value 0.1373) the four chronological groups are very close and share strong genetic affinities compared to other Central and South American samples. Interestingly, initial late Holocene sample is more differentiated from each other and from those of the early and middle Holocene indicating some degree of differentiation.

**Figure 3.**
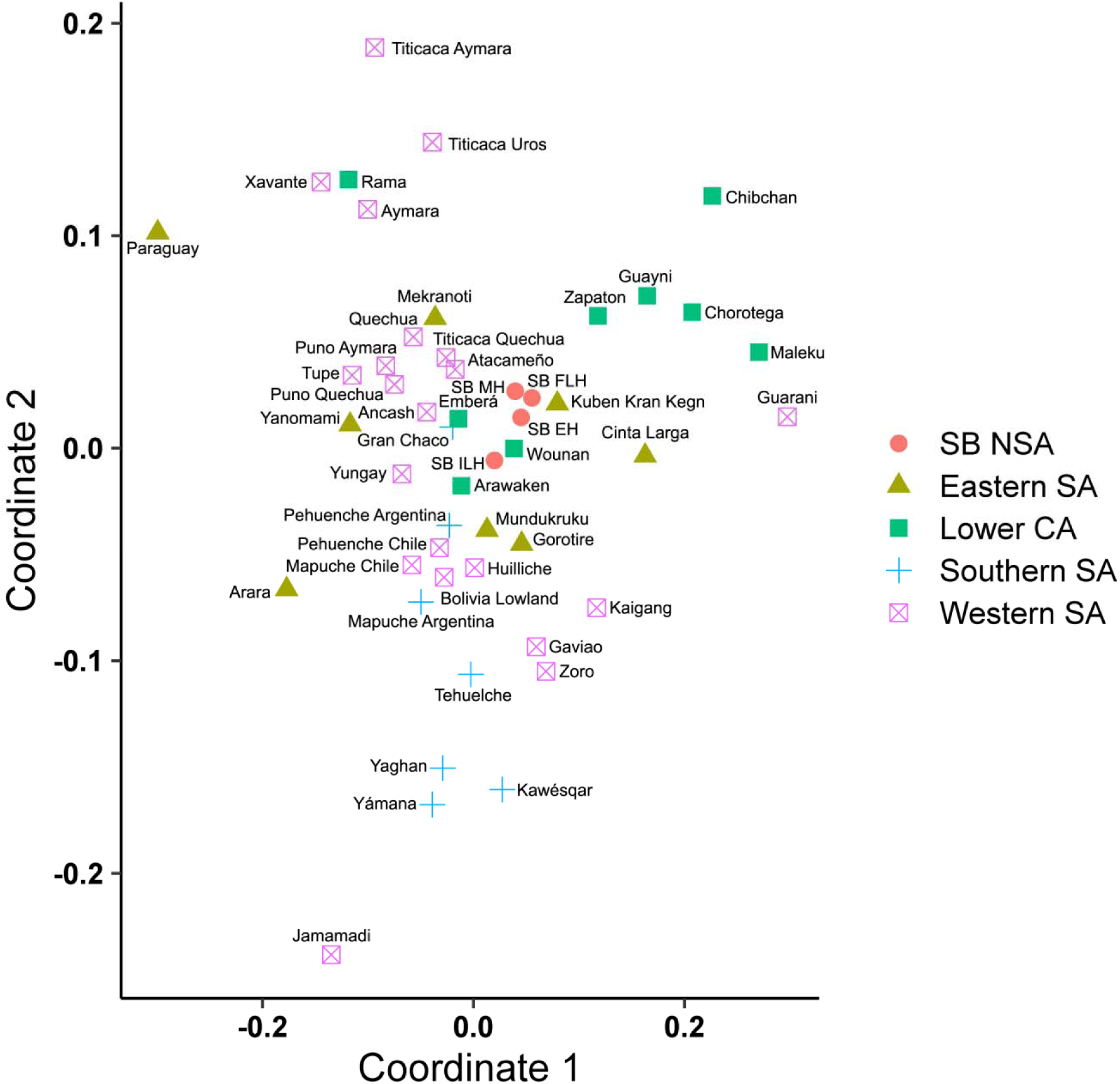
Non-metric multidimensional scaling (MDS) of the inter-population F_ST_ distances for the ancient samples from the Sabana de Bogotá and present-day Native American samples from Central and South America.

**Table 4.** Pairwise inter-population distances (F_ST_, lower left matrix), and p-values (upper right matrix) between the four chronological groups from the Sabana de Bogotá.

|  | EH | MH | ILH | FLH |
| --- | --- | --- | --- | --- |
| EH | 0 | 0.88288+-0.0228 | 0.77477+-0.0279 | 0.78378+-0.0334 |
| MH | -0.10062 | 0 | 0.70270+-0.0466 | 0.72072+-0.0508 |
| ILH | -0.1118 | -0.06949 | 0 | 0.67568+-0.0530 |
| FLH | -0.08741 | -0.05884 | -0.04905 | 0 |

As expected based on the observed F_ST_ inter-population distances between the four chronological groups and the exact test of population differentiation, the demographic simulations with DIYABC returned scenario 1 (population continuity over time) as the most likely scenario with an average posterior probability of 0.8469 (Scenario 2= 0.04; 3= 0.08; 4= 0.03) after 4,000,000 simulations.

## 5. Discussion

Current paleogenomic research with remarkable advances in DNA sequencing technology and computation analysis has revolutionized the study of the Native American population history at regional and continental scales. Important clues on the Native American past have been revealed through the study of ancient genomes, including the early entry of a heterogeneous founding population, its split into different branches, a rapid and structured dispersion south of the Canadás ice sheets likely following a coastal route, and multiple dispersal events from North/Central to South America over the Holocene (Schieb et al., 2018; Posth et al., 2018; Moreno-Mayar et al., 2018).

Despite the SB region having a profuse bioarchaeological record that encompasses the whole Holocene period with a relatively good chronological context, very few reliable paleogenetic studies have been performed to date. This is striking when considering the strategic geographic position of northwest South America for studying the early peopling of the subcontinent. In the present study, we successfully recovered the whole mitogenomes of fifteen individuals dated between ∼8,000 and 4,500 cal BP from the SB region. Furthermore, mtDNA HVR1 sequence data from additional 20 individuals dated between ∼9,000 and 540 cal BP from published studies were included for comparative purposes. The complete dataset analyzed allowed us to investigate the regional population history and, more specifically, to test whether the paleogenetic data supports population continuity or alternatively discontinuity.

### 5.1. Genetic continuity over the Holocene in the Sabana de Bogotá region?

According to the results, the mitochondrial genomes of the four chronological groups investigated fall within the Native American haplogroups A2, B2, C1b, D1 and D4h3a, which are still found among the present-day Native American and admixed populations inhabiting the Eastern Cordillera and neighboring regions in Colombia. We observe a decline of both haplotype (Hd) and nucleotide diversity (π) over time, with lowest values observed in the late Holocene. The pairwise F_ST_ inter-population distances between the four chronological groups indicated no genetic differentiation over time, and the approximate Bayesian Computation (ABC) method indicated that the population continuity over time is the most likely demographic scenario.

Previous studies that also investigated mtDNA data in prehispanic samples from the SB region arrived at similar conclusions (Díaz-Matallana, 2015; Díaz-Matallana et al., 2016). However, in these studies the authors inferred long-standing population continuity between early Holocene and present-day Native Americans and admixed populations, without considering middle, initial late and final late Holocene samples. In addition, these authors misinterpreted the original radiocarbon chronology of the site (Groot, 1992) and assumed that all individuals belong to the earliest occupational events. Direct dating of some of these skeletons confirmed their mostly middle Holocene age (Carvajal et al., 2014; Rodríguez Cuenca in Díaz-Matllana, 2015; Archila et al., 2020). Therefore, such studies cannot be considered as supportive of long-standing population continuity in the SB region over the Holocene but a more restricted one from early to middle Holocene at best (∼9,000-6,000 cal BP). Similarly, studies investigating cranial and dental metric traits have suggested that the remarkable craniodental differentiation viewed throughout time in the SB region is related solely to the effects of local microevolutionary processes (Rodríguez Cuenca, 2007; Rodríguez Cuenca and Vargas, 2010). In their view, neither the arrival of foreign populations(s) (admixing or replacing local populations), nor changes in the subsistence system or environmental conditions promoted large craniofacial and dental transformations/adaptations. These genetic and bioarchaeological studies are relevant in the context of the present mtDNA results supporting population continuity in the SB region. Previously, one of us (Delgado, 2012, 2015, 2017, 2018) has strongly criticized local models of population continuity (LEM) because of their speculative nature, methodological biases and the lack of supporting evidences, and has instead provided evidence favoring the PDM (*contra* Rodríguez Cuenca, 2001, 2007; Rodríguez Cuenca and Vargas, 2010). Nevertheless, in spite of the further support to the PDM from different lines of archaeological and bioarchaeological evidence, this preliminary study supports, to some degree, genetic continuity, or at least not complete replacements and/or admixture over the Holocene in the SB region, which indicates that some variants of the LEM are plausible scenarios. Interestingly, recent studies (Fehren-Schmitz et al., 2015; Llamas et al., 2016; Posth et al., 2018) using high resolution evidence (i.e., whole mitochondrial and nuclear genomes) have revealed that genetic continuity across South America is not an uncommon phenomenon, although it is mostly restricted to some periods (but see Lindo et al., 2018).

A cautionary note is necessary here since on the one hand, we are investigating only the maternal population history, and on the other, we investigated very small sample sizes for all chronological periods, especially for the early Holocene, which translate to a loss of statistical power in the inter-group comparisons. In addition, the data investigated (i.e., HVR1 sequences) would not have the enough resolution to detect subtle but important population changes in complex demographic processes. This implies that the present results must be considered as preliminary and the scenario might change once we generate more data.

### 5.2 Alternative models and hypotheses

Taking into account the previous results arguing against the LEM and the biases of the present study, it seems reasonable to explore alternative explanations for the preliminary results obtained here. According to the PDM, the human history of the SB region cannot be described as a gradual process of *in situ* evolution but on the contrary, a more complex process affected by events of a different nature, such as climatic and environmental shifts, inter-societal contacts, and processes of population contraction, extinction, dispersal, admixture and replacement, as well as dietary adaptations (Delgado, 2012, 2015, 2017, 2018). Here we consider that some of our results would be compatible with the discontinuity scenario. Delgado (2012, 2017) identified at least two periods of major biocultural change, at ∼ 4,700–3,500 cal BP and ∼2,100–1,500 cal BP, that are compatible with several demographic and evolutionary processes. For the first period, non-extensive gene flow was suggested as the likely promoter of the slight cranial diversification observed, whereas for the second period, adaptations to changing economies/diets and genetic drift were proposed as the causal factors of the strong morphological differentiation found. Taking into account the lack of strong genetic differentiation suggested by the mtDNA and the relatively slight cranial diversification observed during the transition from the middle to the initial late Holocene, an assimilation/admixture process between local and extra-regional populations rather than a population replacement or extensive gene flow can account for the cranial and genetic diversity observed. According to the minor morphological and genetic changes inferred, the alleged assimilation/admixture process could have had a relatively minor impact on the genetic makeup of the local populations and the level of resolution used in the present study (i.e., sequences of the HVR1), which makes detecting clear evidences/signals of such a process difficult. The occurrence of an assimilation/admixture process also can explain the persistence of some cultural traits (v.g. the presence of lithic tools using the Abriense technology) and the appearance of new lithic artifacts more related to vegetable processing, new settlement patterns, and atypical mortuary practices during the hunting-gathering to farming transition occurring in the SB region during the middle and initial late Holocene (∼ 6000-4000/3500 cal BP) (Correal, 1990; Delgado, 2012; Triana et al., 2020). Additional studies using more stable (in evolutionary terms) phenotypes than craniometric measurements—such as dental nonmetric traits calculated using quantitative genetic methods (R-matrix)—suggest that minor gene flow during the middle Holocene and genetic drift during the final late Holocene are better explanations for the patterns of dental diversity viewed in the SB region after the second half of the Holocene (Delgado, 2015).

An additional process that can explain the present results under a population discontinuity occurring during the middle to initial late Holocene transition is sex-biased migration. Human population expansions are commonly assumed as involving relatively similar numbers of males and females, but in some cases sex-biased migrations (i.e., involving unequal number of males and females) can also occur, complicating local demographic histories. Previous studies have shown that both short-range and long-range sex-biased migrations are relatively common and respond to different causal factors ranging from cultural practices to climatic and environmental changes (Oota et al., 2001; Langengraber et al., 2007; Rasteiro et al., 2012; Musharoff et al., 2019). Following a highly speculative scenario that takes into account the dental and cranial differentiation suggested in previous studies, it is possible that extra-regional population(s) with a significantly larger number of males arrived to the SB region during the middle to initial late Holocene transition and assimilated (and/or admixed) into local populations. Under this scenario, the local mtDNA diversity remained without extensive changes because the migrating individuals impacting the local gene pool were mostly males.

In addition, short-range migrations (v.g. due to marriage practices) are known to be sex-biased (Oota et al., 2001; Musharoff et al., 2019), thus implying that cultural practices can deeply influence genetic patterns of diversity over time and space. Changes in post-marital residence patterns (PMRP) and the movements of people and genes associated with such practice are thought to be sex-biased (Brewer, 2016; Moravec et al., 2018). For instance, the hunter-gatherer to farmer transition in Europe likely was associated with changes in social organization, particularly in PMRP, indicating that such a practice can lead to different patterns of genetic diversity and differentiation (Rasteiro et al., 2012). Patrilocality and matrilocality are among the most important residence practices influencing human genetic diversity (Oota et al., 2001; Brewer, 2016; Moravec et al., 2018). In the study region, the patterns of post-marital residence have been less studied for the most part of the Holocene and only for the Muisca period some studies have suggested matrilocality (Langebaek, 1989; Correa, 2001). Previous studies supporting population discontinuity (Delgado, 2012, 2015, 2017, 2018) have suggested that extra-regional farmers arrived during the middle/initial late Holocene replacing and/or admixing local hunter-gatherer populations. According to the potential of PMRP to influence within and between-group genetic diversity, sex-biased short-range migrations are an interesting process to be taken into account in future studies modeling SB population history.

Despite additional evidence (v.g., genome wide data) needed to evaluate these scenarios, the hypotheses of an arrival of extra-regional populations with a significantly higher number of males, and/or bias due to post-marital residence practices cannot be rejected here and deserve further evaluation.

### 5.3 Decreasing genetic diversity over time and evidence of gene drift

Two interesting findings of the present research are the decrease in haplotype and nucleotide diversity over time and the evidence of genetic drift and low genetic diversity during the final late Holocene (i.e., samples from the late Muisca period). The linear decrease of mtDNA diversity throughout the Holocene contrasts with the fluctuating cranial diversity observed over time, especially with the high diversity and strong evolutionary diversification seen during the initial late Holocene. The increase in morphological diversity during this period, coincident with the appearance of agriculture, was interpreted as evidence of gene flow (Delgado, 2012, 2015, 2017). The discrepancy between genetics and cranial morphology regarding the amount of diversity deserves attention since a certain degree of convergence would, in principle, be expected from both markers because they are neutral and are affected by similar evolutionary forces (Zichello et al., 2018; von Cramon-Taubadel, 2019). In this specific case, the discrepancy requires additional investigation but, given that some of the cranial phenotypes investigated are very plastic and could reflect a range of adaptations/plasticity rather than exclusively population history (Delgado, submitted), the evolutionary diversification viewed during the initial late Holocene would be likely an effect of natural selection (Delgado, 2017).

Likewise, the diachronic decrease of genetic diversity would suggest both a minor role of gene flow in population differentiation and small population sizes promoting genetic drift (loss of genetic diversity). For the study region, there exist very few studies addressing the role of gene flow and genetic drift in the morphological diversification observed, and the few existing ones have favored the role of gene flow (Delgado, 2012, 2015, 2017; Rodríguez and Colantonio, 2015). On the basis of cemetery size as a proxy of population size (and effective populations size) (see Chamberlain, 2006), Delgado (2015) suggested relatively low population sizes among early and middle Holocene hunter-gatherer populations. Thus, genetic drift likely played a more important role in the reduction of biological diversity and the lack of population differentiation over time at least for some periods. Together these evidences and interpretations reveal that the reconstruction of the local human population histories from integrative approaches is a difficult task that requires new evidences as well as distinct theoretical and methodological approaches.

Apart the evidence of genetic drift and the very low genetic diversity observed during the final late Holocene, this study agrees with previous studies which also found the lowest biological diversity (measured as F_ST_ values) for this period taking into account distinct types of morphological traits, among them cranial shape F_ST_= 0.0143, standard error = 0.003; facial shape F_ST_= 0.006, standard error = 0.003 (Delgado, 2015, 2017); craniofacial allometric shape F_ST_= 0.019, standard error = 0.003 (Delgado, 2012) and dental non-metric traits F_ST_= 0.005, standard error = 0.002 (Delgado, 2015). The very low biological diversity (genetic and morphological) and the likely evidence of genetic drift among the Muisca strongly contrast with its very large population size estimated using camp/settlement and cemetery size (Langebaek, 1995, 2019). In fact, several Muisca cemeteries found around the SB region include hundreds and thousands of burials (Langebaek et al., 2016; Argüello, 2018). According to several archaeological and ethnohistoric studies, the Muisca were among the largest and most socioeconomically complex societies inhabiting the SB region before the arrival of European invaders (Langebaek, 1989; Correa, 2001; Boada, 2007). These data suggest that the Muisca societies presented large population sizes (and effective population sizes), which is incompatible with the action of genetic drift that typically occurs in small populations. Therefore the reduction of biological diversity is likely related to other processes including natural selection and likely endogamy. In the case of morphological diversity, natural selection could be an option. Actually, Delgado (2015, 2017) using quantitative genetic methods (Lande test) proposed that diversifying selection could account for the craniofacial differentiation observed during the final late Holocene. However, the mtDNA is considered a neutral marker and despite the functional importance (v.g., in cellular energy production) of some proteins encoded in the mitochondrial genome, it is not clear how natural selection could have acted in the mtDNA genome of the Muisca populations. Endogamy or inbreeding, on the other hand, is a cultural practice that can produce the reduction of the genetic diversity observed during the final late Holocene, as has been shown in other human and hominin populations (Pemberton et al., 2012; Ríos et al., 2015).

The Muisca sociopolitical organization has been widely studied using ethnohistoric, archaeological and more recently genetic evidences (Broadbent, 1964; Langebaek, 1989, 1995, 2019; Correa, 2001; Kruschek, 2003 Boada, 2007; Pérez L, 2016). In general terms, at the highest level of Muisca sociopolitical organization four main chiefs controlled large regions and received allegiance and tribute from a number of sub-regional and local chiefs of lower rank (Kruschek, 2003). It has been suggested that the Muisca society followed an avunculocal post-marital residence pattern and that the political succession was matrilineal (Villamarin and Villamarin, 1975; Correa, 2001). It is not clear yet if the Muisca practiced full endogamy or exogamy, and/or practiced both depending on the social unit and hierarchy involved. Past ethnohistoric studies have highlighted that endogamy was strongly practiced among the Muisca although it is not clear if this practice can be extended to different social units or is restricted to some of them (Broadbent, 1964). Other ethnohistoric studies, on the other hand, suggested a more exogamous pattern among some kinship groups (Langebaek, 1989). More recently, the few existing genetic studies have suggested that exogamy, polygamy and extensive gene flow can account for the alleged high genetic diversity viewed among Muisca from Tibanica (Pérez L, 2016; Langebaek et al., 2016). Although the social organization of the Muisca society is far from completely understood and complex kinship structure characterized the Muisca chiefdoms at different social and political levels, the strong evidence of remarkable reduction of genetic (and morphological) diversity occurring during this period agrees with past hypotheses of strong endogamy among the Muisca society (Broadbent, 1964) and disagree with the results and interpretations of more recent ethnohistoric and genetic studies (Langebaek, 1989; Correa, 2001; Pérez L, 2016). Interestingly, in societies practicing both patrilocal and matrilocal residence, the local provenance of a couple (i.e., both coming from the same community) promotes endogamy (Brewer, 2016). Thus, endogamy is a process likely involved in the important reduction of genetic diversity seen during the final late Holocene although, as mentioned above, more evidence is required to test this hypothesis.

## 6. Concluding remarks

This study represents one of the most reliable and comprehensive studies to date addressing the Holocene (∼9000–550 cal BP) population history of the Sabana de Bogotá region in Northern South America using paleogenetic evidences. The results obtained support in some degree genetic continuity, or at least not complete replacements and/or admixture, indicating that some variants of the LEM are plausible scenarios. Alternative hypotheses suggest that the present results could support more complex demographic process such as assimilation/admixture and sex-biased migrations. In addition, the linear decrease of mtDNA diversity observed over the temporal sequence investigated could be related to the small population size of early, middle and initial late Holocene populations promoting genetic drift. The remarkable reduction of genetic and morphological diversity inferred for the final late Holocene period could be related to cultural practices such as endogamy and inbreeding rather than genetic drift, given the large effective population sizes among Muisca populations. Discrepancies between genetic and morphological evidences regarding the population continuity/discontinuity can be attributed in part to the confounding effects of natural selection/plasticity acting on some cranial phenotypes. Given the existence of some biases related to small sample sizes, the relatively low resolution of the HVR1 sequences and the fact that we are only studying the maternal side of the local human history, the present study must be considered as preliminary and the local population history scenarios and hypothesis delineated here must be explored using genomic data of higher resolution (v.g., whole mitochondrial and nuclear genomes). Fortunately, such studies are being conducted currently so in the very near future further details on the SB population history will be revealed from paleogenomics.

## Acknowledgments

Special thanks to different institutions for gave us access and permissions to the skeletal collections here investigated including Instituto Colombiano de Antropología e Historia (Bogotá), Museo Nacional de Colombia (Bogotá), Instituto de Ciencias Naturales, Universidad Nacional de Colombia (Bogotá), Universidad de los Andes (Bogotá) and Arge de Colombia S.A (Bogotá). One of us (Freddy Rodríguez) sent to the UCSC Human Paleogenomics Lab (Santa Cruz) the samples here investigated in 2017 and 2018 under the license No 4511 given by Instituto Colombiano de Antropología e Historia in 2014. Arge de Colombia S.A funded part of the laboratory analysis. Thanks to División Antropología, Facultad de Ciencias Naturales y Museo, Universidad Nacional de La Plata (Argentina) and Consejo Nacional de Investigaciones Científicas y Técnicas (CONICET) (Argentina) for the institutional suppor to the first autor (MD).

## References

Aceituno, F.J., Loaiza, N., 2007. Domesticación del bosque en el Cauca Medio colombiano entre el Pleistoceno final y el Holoceno medio. BAR International Series 1654.

Aceituno, F.J, Loaiza, N, Delgado, M.E, Barrientos, G., 2013. The initial human settlement of Northwest South America during the Pleistocene/Holocene transition: Synthesis and perspectives. Quaternary International 301: 23–33.

Andrews, R.M., Kubacka, I., Chinnery, P.F., Lightowlers, R.N., Turnbull, D.M., Howell, N., 1999. Reanalysis and revision of the Cambridge reference sequence for human mitochondrial DNA. Natuare Genetics. 23, 147.

Archila, S., Langebaek, C.H., 2015. Dieta y uso de recursos vegetales de una población humana de hace 5000 años en los Andes Orientales de Colombia-El caso de Ubaté. Informe Final. Instituto Colombiano de Antropología e Historia. Bogotá.

Archila, S., Groot, A.M., Ospina, J.P., Mejía, M., Zorro, C., 2020. Dwelling the hill: traces of increasing sedentism in hunter-gatherers societies at Checua site, Colombia (9500-5052 cal BP). Quaternary International. In press.

Ardila, G., 1984. Chía un sitio precerámico en la Sabana de Bogotá. Fundación de Investigaciones Arqueológicas Nacionales. Banco de la República. Bogotá.

Ardila, G.I, 1991. The peopling of northern South America. En R Bonichsen, KL Turmire (Eds) Clovis: Origins and Adaptations. Center for the Study of the First Americans. Oregon State, Corvallis. p 261–282.

Argüello, P., 2018. Nueva Esperanza 2000 años de historia prehispánica de una comunidad en el Altiplano Cundiboyacense. Xpress Estudio Gráfico y Digital S.A.S. Bogotá.

Bisso-Machado, R., Bortolini, M.C., Salzano, M.F., 2012. Uniparental genetic markers in South Amerindians. Genetics and Molecular Biology, 35: 365–387.

Boada, A.M., 1987. Asentamientos indígenas en el valle de la laguna (Samacá-Boyacá). Fundación de Investigaciones Arqueológicas Nacionales. Banco de la República. Bogotá.

Boada, A.M., 2007. The Evolution of Social Hierarchy in a Muisca Chiefdom of the Northern Andes of Columbia. University of Pittsburgh memoirs in Latin American archaeology no. 17.

Boessenkool, S., Hanghøj, K., Nistelberger, H.M., Der Sarkissian, C., Gondek, A., Orlando, L., Barrett, J.H., Star, B., 2017. Combining bleach and mild pre-digestion improves ancient DNA recovery from bones. Molecular Ecology Resources, 17: 742–751.

Botiva, C.A., 1989. La Altiplanicie Cundiboyasence. In: Botiva, C.A., Cadavid, G., Herrera, L., Groot, A.M., Mora, S., (eds). Colombia prehispánica. Regiones arqueológicas. Bogotá. Universidad Nacional.

Brewer, D., 2016. A systematic review of post-marital residence patterns in prehistoric hunter-gatherers BioRxiv: http://dx.doi.org/10.1101/057059.

Broadbent, S., 1964. Los Chibchas, Organización Sociopolítica. Facultad de Sociología, Universidad Nacional de Colombia, Bogotá.

Bronk Ramsey, C., 2009. Bayesian analysis of radiocarbon dates. Radiocarbon, 51: 337–360.

Cárdenas, F., 2002. Datos sobre la alimentación prehispánica en la Sabana de Bogotá, Colombia. Informes Arqueológicos. Instituto Colombiano de Antropología e Historia. Bogotá.

Carvajal, E., Montes, L., Almanza, O.A. 2014. Datación de restos arqueológicos encontrados en Checua (Cundinamarca-Colombia) mediante resonancia paramagnética electrónica. Revista de la Academia Colombiana de Ciencias, 38: 124–9.

Casas-Vargas, A., Gómez, A., Briceño, I., Díaz, M., Bernal, J., Rodríguez, J.V., 2011. High genetic diversity on a simple of Pre-Columbian bone remains from Guane territories in Northwestern Colombia. American Journal of Physical Anthropology 146: 637–649.

Chamberlain, A.T., 2006. Demography in Archaeology. Cambridge University Press, Cambridge.

Correa, F., 2001. Fundamentos de la organización social Muisca. In: Rodríguez JV(Ed). Los Chibchas. Universidad Nacional de Colombia, 25–48.

Correal, G., 1979. Investigaciones arqueológicas en los abrigos rocosos de Nemocón y Sueva. Fundación de Investigaciones Arqueológicas Nacionales. Banco de la República. Bogotá.

Correal, G., 1986. Apuntes sobre el medio ambiente pleistocénico y el hombre prehistórico en Colombia. In A.L. Bryan (Ed). New Evidence for the Pleistocene Peopling of the Americas. Center for the Study of the Early Man. University of Main, Orono, p 115–131.

Correal, G., 1987. Excavaciones arqueológicas en Mosquera. Revista Estudiantes de Antropología UNAL. 1:13–17.

Correal, G., 1990. Aguazuque: Evidencias de cazadores, recolectores y plantadores en la altiplanicie de la cordillera Oriental. Fundación de Investigaciones Arqueológicas - Nacionales. Banco de la República. Bogotá.

Correal, G., van der Hammen, T., 1977. Investigaciones arqueológicas en los abrigos rocosos del Tequendama. Bogotá. Banco Popular.

Cornuet, J.-M., Pudlo, P., Veyssier, J., Dehne-Garcia, A., Gautier, M., Leblois, R., Marin, J.-M., Estoup, A., 2014. DIYABC v2.0: a software to make approximate Bayesian computation inferences about population history using single nucleotide polymorphism, DNA sequence and microsatellite data. Bioinformatics 30, 1187–1189.

Dabney, J., Knapp, M., Glocke, I., Gansauge, M.-T., Weihmann, A., Nickel, B., Valdiosera, C., García, N., Pääbo, S., Arsuaga, J.-L., Meyer, M., 2013. Complete mitochondrial genome sequence of a Middle Pleistocene cave bear reconstructed from ultrashort DNA fragments. Proceedings of the National Academy of Sciences of the United States of America 110, 15758–63.

Delgado, M.E., 2012. Mid and late Holocene population changes at the Sabana de Bogotá (Northern South America) inferred from skeletal morphology and radiocarbon chronology. Quaternary International, 256:2–11.

Delgado, M.E., 2015. Variación dental y craneofacial en el Norte de los Andes durante el Pleistoceno y el Holoceno. Su relevancia para la discusión de la colonización temprana de Sudamérica, Ph.D. Dissertation. División Antropología, Facultad de Ciencias Naturales y Museo, Universidad Nacional de La Plata, La Plata, Buenos Aires.

Delgado, M.E., 2017. Holocene population history of the Sabana de Bogotá region, Northern South America: an assessment of the craniofacial shape variation. American Journal of Physical Anthropology, 162: 350–369.

Delgado, M.E., 2018. Stable isotope evidence for dietary and cultural change over the Holocene at the Sabana de Bogotá, Northern Andes. Archaeological and Anthropological Sciences, 10: 817–832.

Delgado, M.E., 2020. Craniofacial diversity and the reconstruction of the late Pleistocene and Holocene Native American population history. American Journal of Physical Anthropology. Submitted.

Delgado, M.E., Aceituno, F.J, Loaiza, N. 2015a. Multidisciplinary studies on the human-environment interaction during the initial peopling of the Americas. Quaternary International 363: 1–3.

Delgado, M.E, Aceituno, F.J, Barrientos, G. 2015b. ^14^C data and the early colonization of Northwest South America: a critical assessment. Quaternary International, 363: 55–64.

Días-Matallana, M., 2015. Caracterización genética de un grupo paleoamericano Checua proveniente de Nemocón Cundinamarca, Colombia: implicaciones para el poblamiento temprano de Suarmérica. Ph.D. Dissertation. Pontificia Universidad Javeriana, Facultad de Ciencias, Bogotá.

Días-Matallana, M., Gómez, A., Briceño, I., Rodríguez, J.V., 2016. Genetic analysis of Paleo-Colombians from Nemocón, Cundinamarca provides insights on the early peopling of Northwestern South America. Revista Colombiana de Ciencias Exactas, 40:461–483.

Dillehay, T.D., 2000. The settlement of the Americas. A new prehistory. Basic Books.

Dueñas, H., 1980. Variaciones climáticas del Pleistoceno superior y del Holoceno en la Sabana de Bogotá (estudio palinológico de la formación sabana). Antropológicas 2, 31–38.

Enciso, B., 1990-1991. Arqueología de rescate en el Barrio Las Delicias (Bogotá). Revista Colombiana de Antropología, 28:57–60.

Excoffier, L., Lischer, H.E., 2010. Arlequin suite ver 3.5: a new series of programs to perform population genetics analyses under Linux and Windows. Molecular Ecology Resources, 10: 564–567.

Fehren-Schmitz, L., Llamas, B., Lindauer, S., Tomasto-Cagigao, E., Kuzminsky, S., Rohland, N., et al. 2015. A re-appraisal of the early Andean human remains from Lauricocha in Peru. Plos One, 10: e0127141.

Gómez, A., Berrío, J., Hooghiemstra, H., Becerra, M., Marchant, R., 2007. A Holocene pollen record of vegetation change and impact from Pantano de Vargas, and Intra-Andean basin of Duitama, Colombia. Review of Paleobotany and Palynology, 145, 143–157.

Goudet, J., Raymond, M., Meeus, T. de, Rousset, F., 1996. Testing differentiation in diploid populations. Genetics 144, 1933–1940.

Gnecco, C., 1999. An archaeological perspective of the Pleistocene/Holocene boundary in northern South America. Quaternary International, 53/54: 3–9.

Gnecco, C., 2003. Against ecological reductionism: Late Pleistocene hunter-gatherers in the tropical forests of northern South America. Quaternary International, 109-110:13-21.

Groot, A.M., 1992. Checua. Una secuencia cultural entre 8500 y 3000 años antes del presente. Fundación de Investigaciones Arqueológicas Nacionales. Bogotá.

Hogg, A. G., Hua, Q., Blackwell, P. G., Niu, M., Buck, C. E., Guilderson, T. P., Heaton, T. J., Palmer, J. G., Reimer, P. J., Reimer, R. W., Turney, C. S. M., & Zimmerman, S. R. H., 2013. SHCal13 Southern Hemisphere Calibration, 0-50,000 Years cal BP. Radiocarbon, 55: 1889-103.

Hurt, W.R., van der Hammen, T., Correal, G., 1977. The El Abra rock shelters, Sabana de Bogotá, Colombia, South America. Occasional papers and monographs, vol 2. Bloomignton Indiana University Museum.

Jónsson, H., Ginolhac, A., Schubert, M., Johnson, P.L.F., Orlando, L., 2013. MapDamage2.0: Fast approximate Bayesian estimates of ancient DNA damage parameters. Bioinformatics, 29: 1682–1684.

Kruschek, M.H., 2003 The Evolution of the Bogotá Chiefdom: a Household View. Ph.D Dissertation. Faculty of Arts and Sciences. Department of Anthropology, University of Pittsburgh.

Langebaek, C.H., 1989. Mercados, poblamiento e integración étnica entre los Muiscas del siglo XVI. Banco de la República, Bogotá, Colombia.

Langebaek C.H., 1995. Arqueología regional en el territorio Muisca. Estudio de los valles de Fúquene y Susa. University of Pittsburg. Department of Anthropology. Memoirs in Latin American Archaeology. No 9.

Langebaek C.H., 2019. Los muiscas: la historia milenaria de un pueblo chibcha. Editorial Debate, Bogotá, Colombia.

Langebaek C.H., Jaramillo, A., Aristizábal, L., Bernal, M., Corcione, M.A., Mendoza, L., Pérez, L., Rodríguez, F., Zorro, C., 2016. Vivir y morir en Tibanica: reflexiones sobre el poder y el espacio en una aldea Muisca tardía de la Sabana de Bogotá. Revista Colombiana de Antropología, 51:173–207.

Langergraber, K.E., Siedel, H., Mitani, J.C., Wrangham, R.W., Reynolds, V., Hunt, K., et al., 2007. The genetic signature of sex-biased migration in patrilocal chimpanzees and humans. Plos One 2, e973.

Lindo, J., Haas, R., Hoffman, C., Apata, M., Moraga, M., Verdugo, R., 2018. The genetic prehistory of the Andean highlands 7000 years BP though European contact. Sciences Advances 4: eaau4921.

López, C., 2008. Landscape Development and the Evidence for Early Human Occupation in the Inter-Andean Tropical Lowlands of the Magdalena River, Colombia. Syllaba Press, Miami.

Li, H., Durbin, R., 2009. Fast and accurate short read alignment with Burrows-Wheeler transform. Bioinformatics, 25: 1754–60

Llamas, B., Fehren-Schmitz, L., Valverde, G., Soubrier, J., Mallick, S., Rohland, N., et al., 2016. Ancient mitochondrial DNA provides high-resolution time scale of the peopling of the Americas. Sciences Advances, 2: e1501385.

Llamas, B., Valverde, G., Fehren-Schmitz, L., Weyrich, L.S., Cooper, A., Haak, W., 2017. From the field to the laboratory: Controlling DNA contamination in ancient DNA research. STAR-Science and Technology in Archaeological Research 3, 1–14.

Marchant, R., Behling, H., Berrío, J., Cleef, A., Duivenvoorden, J., Hooghiemstra, H., Kuhry, P., Melief, R., Schreve-Brinkman, E., van Geel, B., van der Hammen, T., van Reenen, G., Wille, M., 2002. Pollen-based biome reconstructions for Colombia at 3000, 6000, 9000, 12000, 15000 and 18000 ^14^C yr ago: late Quaternary tropical vegetation dynamics. Journal of Quaternary Science, 17: 113-129.

Melton, P.E., Baldi, N.F., Barrantes, R., Crawford, M.H., 2013. Microevolution, migration, and the population structure of five Amerindian populations from Nicaragua and Costa Rica. American Journal of Human Biology, 25:480–490.

Meyer, S., Weiss, G., Haeseler, A. von, 1999. Pattern of nucleotide substitution and rate heterogeneity in the hypervariable regions I and II of human mtDNA. Genetics, 152: 1103– 1110.

Miller, M., 2016. Social Inequality and the body: diet, activity, and health differences in a prehistoric Muisca population (Sabana de Bogotá, Colombia, AD 1000-1400). Ph.D. Dissertation. University of California, Berkeley.

Moravec, J., Atkinson, Q., Bowern, C., Greenhill, S., Jordan, F., Ross, R., et al., 2018. Post-marital residence patterns show lineage-specific evolution. Evolution and Human Behavior, 39: 594–601.

Moreno-Mayar, J.V., Vinner, L., de Barros Damgaard, P., de la Fuente, C., Chan, J., Spence, J.P., et al. 2018. Early human dispersals within the Americas Science, 362: eaav2621

Musharoff, S., Shringarpure, S., Bustamante, C.D., Ramachandran, S., 2019. The inference of sex-biased human demography from whole-genome data. Plos Genetics, 15: e1008293

Muttillo, B., Lembo, G., Rufo, E., Peretto, C., Pérez, R.L., 2017. Revisiting the oldest known lithic assemblages of Colombia: a review of data from El Abra and Tibitó (Cundiboyacense Plateau, Eastern Cordillera, Colombia). Journal of Archaeological Sciences Reports, 13: 455–465.

Neves, W.A., Hubbe, M., Correal, G., 2007. Human skeletal remains from Sabana de Bogotá, Colonbia: A case of Paleoamerican morphology late survival in South America? American Journal of Physical Anthropology, 130: 1080–1098.

Nieuwenhuis, C.J., 2002. Traces on tropical tools: a functional study of chert artifacts from preceramic sites in Colombia. PhD Thesis from Leiden University. Archaeological Studies Leiden University No. 9. Faculty of Archaeology, University of Leiden, Leiden.

Oota, H., Settheetham-Ishida, W., Tiwawech, D., Ishida, T., Stoneking, M., 2001. Human mtDNA and Y-chromosome variation is correlated with matrilocal versus patrilocal residence. Nature Genetics, 29:20–21.

Orrantia, J.C., 1997. Potreroalto: informe preliminar sobre un sitio temprano en la Sabana de Bogotá. Revista de Antropología y Arqueología, 9: 181–184.

Ospina, J.P., Archila, S., 2020. Marking graves and intruding on the dead: An archaeothanatological analysis to unveil posthumous experiences of death and remembrance at the site of Checua, Colombia (7580-5052 cal BP). Quaternary International. In press.

Oven, M. van, 2015. PhyloTree Build 17: Growing the human mitochondrial DNA tree. Forensic Science International: Genetics Supplement Series, 5: e392–e394.

Oven, M. van, Kayser, M., 2009. Updated comprehensive phylogenetic tree of global human mitochondrial DNA variation. Human Mutation, 30: E386–E394.

Pemberton, T., Fang-Yuan, L., Hanson, E., Mehta, N., Choi, S., Ballantyne, J, 2012. Impact of restricted Marital Practices on Genetic Variation in an Endogamous Gujarati Group. American Journal of Physical Anthropology, 149: 92–103.

Pérez, L., 2015. Aportes genéticos para el entendimiento de la organización social de la comunidad Muisca Tibanica (Soacha, Cundinamarca). Ph.D. Dissertation, Departamento de Ciencias Biológicas, Universidad de los Andes, Bogotá, Colombia.

Pérez, L., Rodríguez, F., Langebaek, C., Groot, H., 2016. Cuantificación en tiempo real de un conjunto de muestras colombianas de relevancia histórica mediante un fragmento corto de la región hipervariable II del ADN mitocondrial. Biomédica, 36:475–482.

Pérez, P., 2001. Procesos de interacción en el área septentrional del altiplano Cundiboyacense y oriente de Santander. In Rodríguez JV(ed). Los Chibchas. Universidad Nacional de Colombia, 49–110.

Pinto, M., 2003. Galindo, un Sitio a Cielo Abierto de Cazadores-Recolectores en la Sabana de Bogotá. Fundación de Investigaciones Arqueológicas Nacionales, Banco de la República, Bogotá.

Posth, C., Nakatsuka, N., Lazaridis, L., Skoglund, P., Mallick, S., Lamnidis, T., et al 2018. Reconstructing the deep population history of Central and South America. Cell, 175: 1185–1197.

Rasteiro, R., Boutier, P.A., Souza, V., Chikhi, L., 2012. Investigating sex-biased migration during the Neolithic transition in Europe, using an explicit spatial simulation framework. Proceedings of the Royal Society B, 279: 2409–2416.

Renaud, G., Slon, V., Duggan, A.T., Kelso, J., 2015. Schmutzi: estimation of contamination and endogenous mitochondrial consensus calling for ancient DNA. Genome Biology 16, 224.

Ríos, L., Rosas, A., Estalrrich, A., García-Tabernero, A., Bastir, M., Huguet, R., et al. 2015. Possible Further Evidence of Low Genetic Diversity in the El Sidrón (Asturias, Spain) Neandertal Group: Congenital Clefts of the Atlas. Plos One, 10: e013655.

Rodríguez Cuenca, J.V., 2001. Craneometría de la población prehispánica de los Andes Orientales de Colombia: diversidad, adaptación y etnogénesis. Implicaciones para el poblamiento Americano. In: Rodríguez JV(ed). Los Chibchas. Universidad Nacional de Colombia, 251–310.

Rodríguez Cuenca, J.V., 2007. La diversidad poblacional de Colombia en el tiempo y en el espacio: estudio craneométrico. Revista de la Academia Colombiana de Ciencias. Vol XXXI (120): 321-346.

Rodríguez Cuenca, J.V., Vargas, C., 2010. Evolución y tamaño dental en poblaciones humanas de Colombia. Revista de la Academia Colombiana de Ciencias 34: 423–439.

Rodríguez Flórez, C.D., Colantonio, S., 2015. Biological affinities and regional micro-evolution among prehispanic communities of Colombiás northern Andes. Anthropologischer Anzeiger, 72: 141–168.

Rohland, N., Harney, E., Mallick, S., Nordenfelt, S., Reich, D., 2015. Partial Uracil -DNA-treatment for screening of ancient DNA. Philosophical Transactions of the Royal Society of London B, 370:20130624.

Scheib, C.L., Li, H., Desai, T., Link, V., Kendall, C., Dewar, G., et al 2018. Ancient human parallel lineages within North America contributed to a coastal expansion. Science, 360:1024–1027.

Skoglund, P., Stor\a a, J., Götherström, A., Jakobsson, M., 2013. Accurate sex identification of ancient human remains using DNA shotgun sequencing. Journal of Archaeological Science, 40: 4477–4482.

Slatkin, M., 1995. A measure of population subdivision based on microsatellite allele frequencies. Genetics, 139: 457–462.

Soares, P., Ermini, L., Thomson, N., Mormina, M., Rito, T., Röhl, A., Salas, A., Oppenheimer, S., Macaulay, V., Richards, M.B., 2009. Correcting for purifying selection: An improved human mitochondrial molecular clock. American Journal of Human Genetics, 84: 740–759.

Triana, A., 2019. Dieta y acceso a recursos determinados a partir del sexo en grupos cazadores-recolectores del Holoceno temprano y medio de la Sabana de Bogotá. Ph.D. Dissertation. Departamento de Antropología, Facultad de Ciencias Sociales, Universidad de los Andes, Bogotá.

Triana, A., Casar-Aldrete, I., Morales-Puente, P., Salinas-Acero, J., 2020. Isótopos estables en restos óseos humanos y de fauna, sitios arqueológicos de Tequendama y Aguazuque, Holoceno temprano y medio sabana de Bogotá, Colombia. Jangwa Pana. In press.

Tamura, K., Nei, M., 1993. Estimation of the number of nucleotide substitutions in the control region of mitochondrial DNA in humans and chimpanzees. Mol.Biol.Evol. 10, 512– 526.

Troll, C.J., Kapp, J., Rao, V., Harkins, K.M., Cole, C., Naughton, C., Morgan, J.M., Shapiro, B., Green, R.E., 2019. A ligation-based single-stranded library preparation method to analyze cell-free DNA and synthetic oligos. BMC Genomics, 20: 1023.

van der Hammen, T., 1974. The Pleistocene changes of vegetation and climate in tropical South America. Journal of Biogeography 1: 3–26.

van der Hammen, T., González, E., 1960. Upper Pleistocene and Holocene climate and vegetation of the Sabana de Bogotá (Colombia, South America). Laidse Geologische Mededelingen 25, 261–315.

van der Hammen, T., Correal, G., 1992. El hombre prehistórico en la Sabana de Bogotá: datos para una prehistoria ecológica. In: van der Hammen, T (ed). Historia, ecología y vegetación. Corporación Colombiana para la Amazonia. Araraucara (Bogotá), 211–230.

van der Hammen, T., Correal, G., van Klinken, G.J., 1990. Isótopos estables y dieta del hombre prehistórico en la Sabana de Bogotá. Boletín de Arqueología. Fundación de Investigaciones Arqueológicas Nacionales, 5: 3–10.

van der Hammen, T., Hooghiemstra, H., 1995 The El Abra stadial, a Younger Dryas equivalent in Colombia. Quaternary Sciences Review. 14, 841–851.

Vélez, M., Hooghiemstra, H., Metcalfe, S., Willie, M., Berrío, J., 2006 Late Glacial and Holocene environmental and climatic changes from a limnological transects through Colombia, northern South America. Palaeogeography, Palaeoclimatology, Palaeoecology. 234, 81–96.

Villamarin, J.A., Villamarin, J.E., 1975. Kinship and inheritance among the Sabana de Bogotá Chibcha at the time of Spanish conquest. Ethnology, 14: 173–179.

Von Cramon-Taubadel, N., 2019. Multivariate morphometrics, quantitative genetics, and neutral theory: Developing a “modern synthesis” for primate evolutionary morphology. Evolutionary Anthropology, 28: 21–33.

Wang, S., Lewis, C., Jakobsson, M., Ramachadran, S., Ray, N., Bedoya, G., et al. 2007. Genetic variation and population structure in Native Americans. Plos Genetics 3(11): e185.

Weissensteiner, H., Pacher, D., Kloss-Brandstätter, A., Forer, L., Specht, G., Bandelt, H.-J., Kronenberg, F., Salas, A., Schönherr, S., 2016. HaploGrep 2: mitochondrial haplogroup classification in the era of high-throughput sequencing. Nucleic Acids Research 44, W58– W63.

Zichello, J., Baab, K., McNulty, K., Raxworthy, C., Steiper, C., 2018. Hominoid intraspecific cranial variation mirrors neutral genetic diversity. Proceedings of the National Academy of Sciences, 115: 11501–11506.

